# A new genome-wide method to identify genes with bimodal gene expression

**DOI:** 10.1101/2020.12.21.423759

**Authors:** Josivan Ribeiro Justino, Clovis F. Reis, Andre Faustino Fonseca, Sandro Jose de Souza, Beatriz Stransky

## Abstract

A new method is presented to detect bimodality in gene expression data using the Gaussian Mixture Models to cluster samples in each mode. We have used the method to search for bimodal genes in data from 25 tumor types available from The Cancer Genome Atlas. The method identified 554 genes with bimodal gene expression, of which 46 were identified in more than one cancer type. To further illustrate the impact of the method, we show that 96 out of the 554 genes with bimodal expression patterns presented different prognosis when patients belonging to the two expression peaks are compared. The software to execute the method and the corresponding documentation are available at https://github.com/LabBiosystemUFRN/Bimodality_Genes.

## Introduction

Studies on gene expression and regulation have been directed towards a better understanding of a diverse range of biological processes, including the initial differentiation in the embryonic stage and changes in health and disease that occur during life. These patterns of gene expression have been extensively used to establish associations between phenotypes and genetic/epigenetic information [1–2]. The challenges for such studies are significant, however, and the identification of expression signatures enriched with bona fide phenotypic associations is particularly welcome. In that aspect, bimodal gene expression is an interesting pattern since their identification capitalizes on the availability of genetic and clinical data from large cohorts of samples and each mode can, in theory, correspond to a phenotypic state of the system.

Few previous studies have searched for bimodality in large-scale gene expression data [3–5]. Causes for such bimodality have been discussed, including: i) differential action of transcription factors [6], ii) regulation by microRNAs [7, 8]; iii) regulation by circular RNA [9] and even iv) stochastic events [10].

The identification of genes with a bimodal expression pattern, together with sample stratification, can be used to identify important clinical and therapeutic targets [11]. Furthermore, this process can reveal molecular signatures that distinguish tumor subtypes, which would contribute to a better clinical understanding of the biological characteristics of cancer. To be clinically useful, a bimodal pattern must exhibit a clear separation between the two groups and have significant sample sizes [12]. The term “bimodal expression” is related in biology to two distinct groups of continuous values of gene expression for the same gene. Statistically, the set of continuous values of lower and higher expression has a more consistent definition as a mixture of Gaussian distributions.

Here, a computational protocol was developed to identify genes with bimodal expression patterns using the Gaussian Mixture Models (GMM) in a genome-wide context. To prove the applicability and robustness of our method, we used this new tool to identify genes with bimodal expression in 25 tumor types whose expression data is available from The Cancer Genome Atlas (TCGA). Finally, we made use of the availability of clinical data from TCGA to find 96 genes, among the ones with bimodal gene expression, in which patients in the two expression peaks showed different prognosis. Software to execute the method and the corresponding documentation are freely available at https://github.com/LabBiosystemUFRN/Bimodality_Genes.

## Materials & Methods

### Data samples

Expression and clinical data from 25 different tumor types were obtained from The Cancer Genome Atlas (TCGA) project through the Genomic Data Commons Data Portal (https://portal.gdc.cancer.gov). Expression data for 24,456 genes were evaluated to identify genes with a bimodal distribution, using Fragments by Exon Kilobase per Millions of Mapped Fragments (FPKM) values. For survival analysis, the clinical information was extracted from cBioPortal for Cancer Genomics (https://www.cbioportal.org/).

### Detection of bimodality

The detection of bimodality involves a three-step process, configured by 7 parameters, listed below:

a. minExpression - defines the minimum expression value in the analysis. It prevents noise in readings of low expression value from influencing the correct detection of peaks, particularly at values close to zero. This parameter must be appropriate to the type of measurement unit of expression to be used. Its default value is 0.02 FPKM;
b. minSampleSize - defines the minimum sample size in the analysis. Datasets with a number of samples smaller than this value do not undergo any processing, being immediately discarded. Its default value is 50 samples.
c. MinClusterSize - defines the minimum size, in relation to the total number of samples, that a cluster must have to be considered as one of the bimodal clusters. This aims to discard groups of relatively small sample populations composed of outliers, capable of altering the density profile to the point of being mistaken as a peak, especially when they occur in the upper tail of the distribution. Its default value is 10% of the total samples considered.
d. Threshold Up - defines the minimum difference between the points detected as adjacent peaks and valleys on the density curve. If the difference between them is less than the parameter value, this oscillation in the density graph will be disregarded in the detection process. Its default value is 10% of the maximum density value.
e. Threshold Down - peaks whose density values are below this threshold will be discarded. This aims to rule out small fluctuations in the expression values that normally occur in the upper tail of the distribution, which cause the density to fluctuate widely in this region. Its default value is 20% of the maximum density value.
f. Smoothing factor – this parameter mitigates the variations in the derivatives curve to make detection less sensitive. Its default value is ‘true’.
g. useLog – this parameter defines whether the expression values will be considered in their original form or whether they should be transformed into a base 2 or base 10 logarithm before analysis. This helps to improve the sensitivity of the algorithm, particularly when the range of expression values is quite wide, which causes the upper tail of the density curve to flatten, making the peak detection process more difficult. An example of this difference in the density profile can be seen in Supplementary Figure 1 where the same dataset has its density curve plotted with and without the log_10_ transformation of the expression values. Its default value is “none”.

The three steps are:

a. Peak detection: In this step, the initial screening of candidate genes for bimodality is performed using the density derivative. First, the density of the expression distribution of each gene is calculated using the density function of the R stats package [13], with the “nrd0” method to calculate the smoothing bandwidth. This method was chosen specifically because it is less precise than methods like the Sheater Jones bandwidth, guaranteeing only the detection of large fluctuations in density. Next, the first density derivative is calculated, which undergoes a smoothing process designed to decrease the sensitivity of peak detection. For this purpose, the smooth.spline function of the R stats package was used [13], with the parameter defined by the smoothing factor. Derivative values tending to zero indicate a peak or valley. The threshold Up and Down parameters are then applied, which will define which peaks can be considered relevant. As a result, this process returns the estimated number of peaks, which will become variable k in the subsequent step.
b. Clustering A data model that presents a characteristic of bimodality can be considered as the overlap of probabilistic models that represent two distinct subpopulations. In this way, we can consider bimodal distributions as a model of mixing Gaussian data (Gaussian Mixture Models - GMM) and use their specific algorithms to perform the identification and separation of these subpopulations [14]. To perform the classification based on GMM, we used the Mclust function of the R mclust package [15], which performs a clustering of data using the expectation maximization (EM) technique, performing successive grouping operations and comparing groups with a Gaussian distribution [16,17]. This process can either infer the number of clusters expected in the distribution or start from a ‘k’ parameter that will designate the number of desired clusters. In our case, we already have such information, the number of clusters will be equal to the number of peaks, estimated in the previous step, plus one. Consequently, the algorithm will group the data in k effective clusters, plus an additional cluster (k + 1), that will contain all data points with low affinity to the main clusters. This process returns, in addition to the clustering of samples in k + 1 clusters, an uncertainty index related to such classification. Arbitrarily, only the samples whose reliability in the classification received an index higher than 46% are maintained. This low rigor in the use of uncertainty values is justified because a distribution of expression indexes tends to be closer to a Poisson than to a Gaussian [18–20], and an excessive rigor in the use of such reliability would cause large disposal of samples; The result of the process is illustrated in Supplementary Figure 2. After discarding clusters smaller than MinClusterSize (m) and samples contained in cluster k + 1, the remaining samples are passed on to the third phase of the process.
c. Peak confirmation The samples contained in the largest k-m clusters are subjected to a new peak detection process, identical to step 1, to confirm the initial screening. If the Peak Detection process continues to identify a bimodality, as is the case shown in Supplementary Figure 3A, that gene is classified as bimodal. Otherwise, the gene is discarded from the bimodal gene pool. Such a situation can be seen in Supplementary Figure 3B, where the bimodality existing in the original dataset no longer can be identified when the filtered samples are used.

### Survival Analysis

To verify if individuals belonging to the two different peaks of expression in a bimodal gene presented a significant difference in survival curves, we performed an analysis using the clinical data from CBioPortal, obtained as indicated above, and the Survival package in R [21].

The samples identified as peak 1 and peak 2 from 554 genes with bimodal distribution were selected and Kaplan-Meier curves were evaluated with a significance level of 5% and 1% using the log-rank test. Kaplan-Meier curves were plotted using the ggplot2 package [22].

## Results

### Development of a method to identify genes with bimodality in gene expression data

In our method, described in Figure 1A, the identification of gene expression bimodality involves a density function, which can be used to analyze the expression values (in FPKM) for all human genes in any set of samples. Using a computational algorithm in R, the maximum and minimum points in the expression density curve of each gene is defined by identifying where the values of its derivative curve change its value signal (Figure 1B). To avoid possible noises in the stratification of samples, data points below 0.002 FPKM were excluded. To identify robust distributions concerning the difference in bimodality peaks, two thresholds were established: (i) a maximum value of 10% of density, used specifically to eliminate small ripples in the upper tail of the distribution curve, which could indicate irrelevant peaks; (ii) a 5% difference between the peak and the density valley, guaranteeing significant differences in the bimodality peaks. All these parameters are shown in Figure 1B in a schematic bimodal distribution of a hypothetical gene.

**Fig 1.**
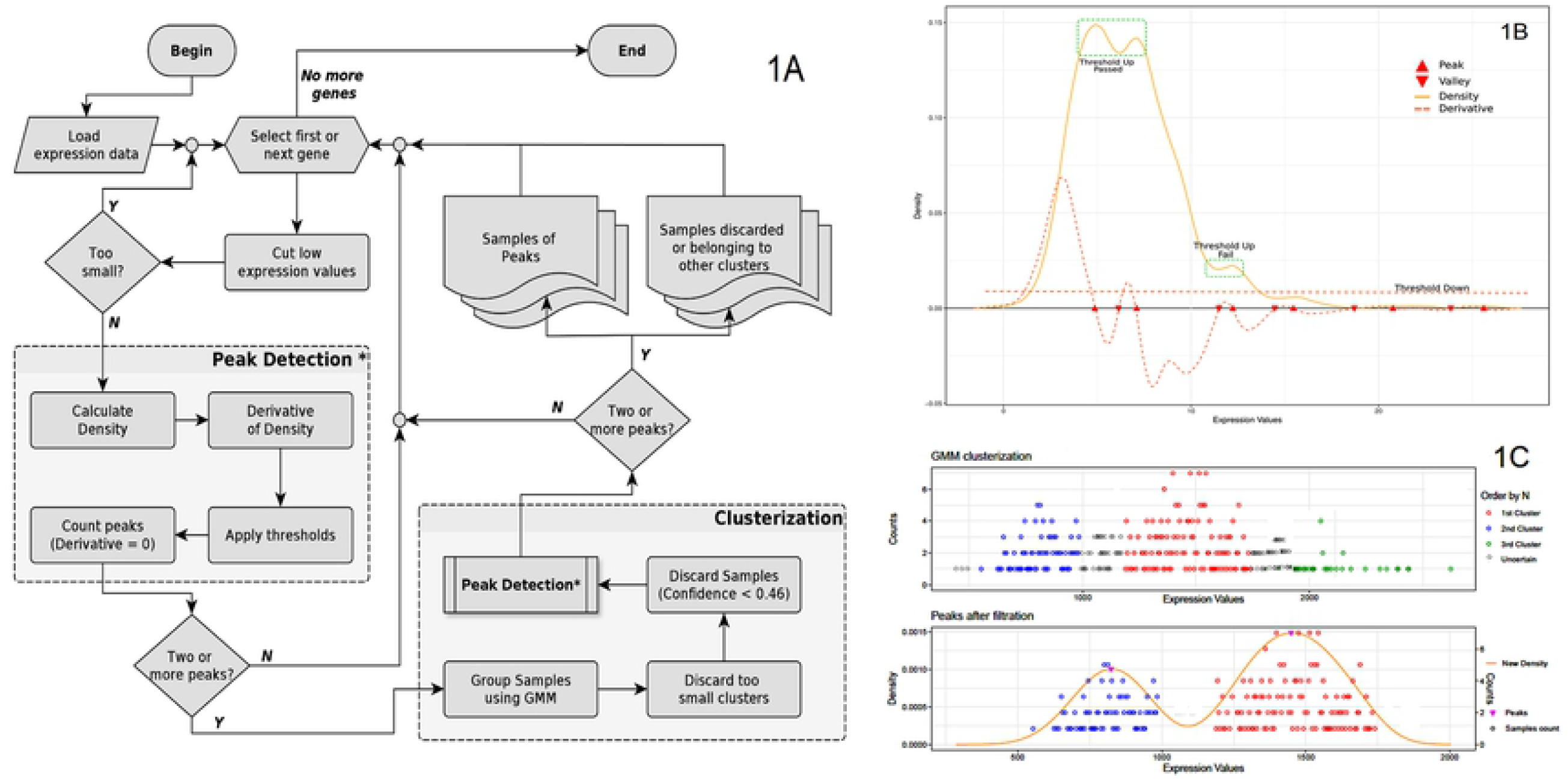
Computational scheme for the identification of genes showing bimodal gene expression patterns. 1A) Stages performed to process the data. 1B) Schematic view of a hypothetical gene with bimodal expression with all important parameters used to define bimodality indicated. 1C) Schematic view of sample clustering process, which identifies samples belonging to each peak in the bimodal distribution (see main text for more details).

After the identification of genes with a bimodal expression pattern, we next performed sample stratification to identify samples belonging to the first and second peaks. Assuming that each peak of the density curve represents the fashion of a subpopulation with different expression features, we can consider that the distribution of expression values, of genes with bimodality characteristics, constitutes a mixture model. Therefore, to ensure the reliable selection of samples according to the selected bimodal genes, an analysis based on the Gaussian Mixture Models (GMM) probabilistic model was introduced in our computational protocol. A schematic view of the sample stratification step is shown in Figure 1C. Finally, stratified samples were submitted to the step of bimodality detection again. Only genes that remained with a bimodal pattern after sample stratification are listed in our final results.

### Identification of bimodal genes using data from 25 different tumor types from TCGA

To illustrate the use of our method, we have collected all gene expression and clinical data from the TCGA for the following tumors: BLCA, BRCA, CESC, COADREAD, ESCA, GBM, HNSC, KIRC, KIRP, LAML, LGG, LIHC, LUAD, LUSC, OV, PAAD, PCPG, PRAD, SARC, SKCM, STAD, TGCT, THCA, THYM, UCEC. These tumors were selected because they have a minimum number of 100 patients in their dataset. A total of 554 unique genes was identified as having a bimodal pattern for at least one tumor type (analysis performed without the logarithmic transformation). Table 1 shows the numbers of genes identified as bimodal for each tumor type (listed at Supplementary Table 1) and Figure 2 shows the bimodal pattern of expression for 25 genes, arbitrarily selected, one for each tumor type. Supplementary Figure 4 shows the same type of plot for all genes found to have a bimodal expression pattern.

**Figure 2:**
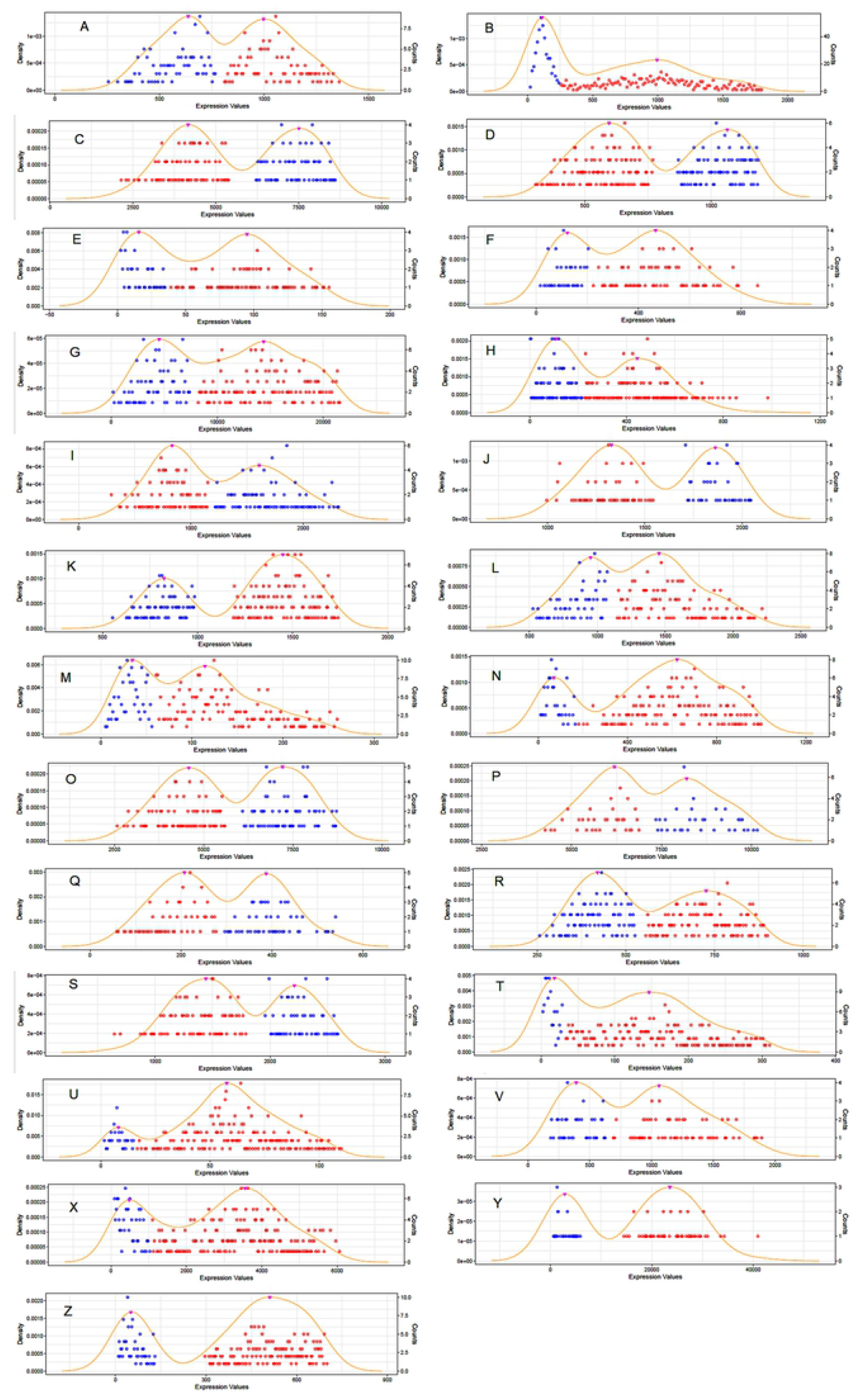
Expression plot showing bimodality for a selection of genes. A) Gene LRRC14 for BLCA; B) Gene RPS27 for BRCA; C) Gene SEP2 for CESC; D) Gene CHMP7 for COADREAD; E) Gene ZNF502 for ESCA; F) Gene GPX8 for GBM; G) Gene CDH3 for HNSC; H) Gene UTY for KIRC; I) Gene FUK for KIRP; J) Gene ADSL for LAML; K) Gene FOXJ3 for LGG; L) Gene ALG8 for LIHC; M) Gene TMLHE for LUAD; N) Gene MTAP for LUSC; O) Gene RCC2 for OV; P) Gene CD164 for PAAD; Q) Gene GMIP for PCPG; R) Gene XRRA1 for PRAD; S) Gene EI24 for SARC; T) Gene PPAPDC3 for SKCM; U) Gene ZNF597 for STAD; V) Gene PCMTD1 for TGCT; X) Gene PLCD3 for THCA; Y) Gene ARHGDIB for THYM and Z) Gene MLH1 for UCEC.

**Table 1.**
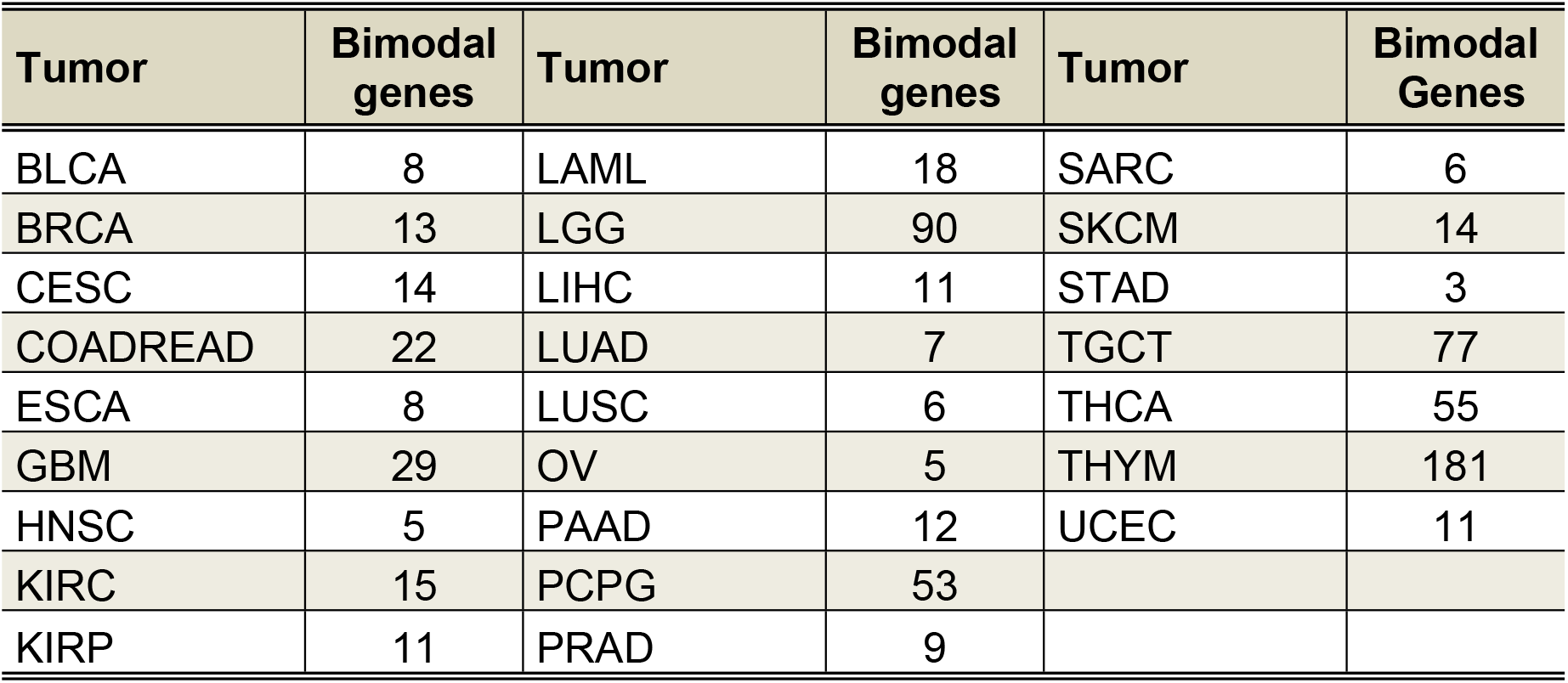
Number of genes showing bimodality for each tumor type.

We found 46 genes showing bimodal expression in more than one tumor type (Table 2). The ones most frequently found were SLC35E2, EIF1AY and RPS27, which have a bimodal pattern in 19, 10 and 10 tumor types, respectively. In this list of genes, chromosomal distribution is significantly biased toward the Y chromosome (p<10^-5^), as observed in Table 2.

**Table 2:**
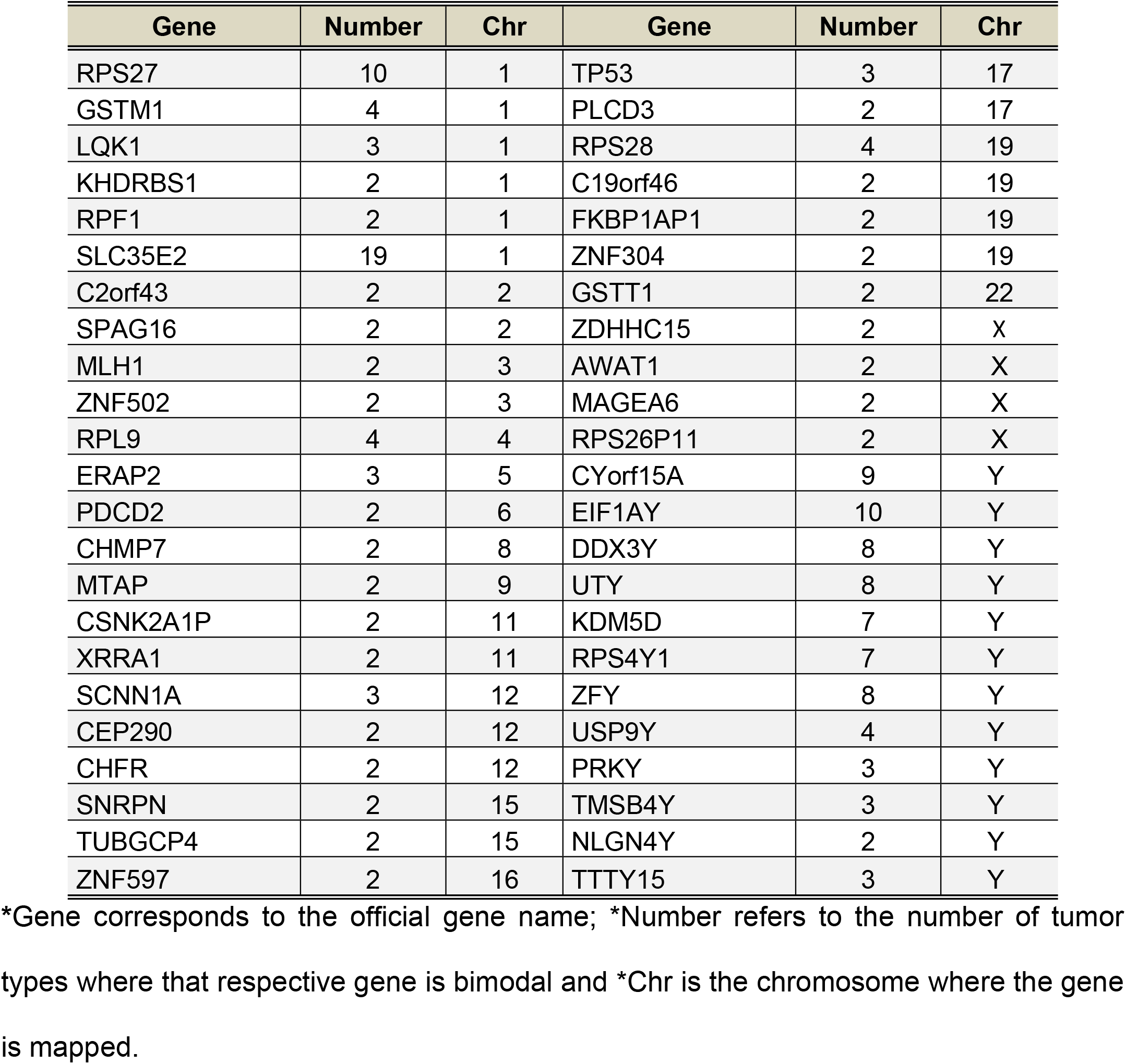
List of genes showing bimodality for more than one tumor type.

### Patients in different expression peaks have different prognosis

We wondered whether patients belonging to the two different peaks of the bimodal distribution would present different prognosis, as evaluated by survival curves in a Kaplan-Meyer plot. All genes identified as having a bimodal distribution (Table 1) were tested. A total of 96 genes were identified as having their bimodal pattern significantly (p<0.01) associated with prognosis (samples belonging to the first peak having either a better or worse prognosis when compared to samples belonging to the second peak). If a threshold of p<0.05 is used, 176 genes are identified as associated with prognosis.

Figure 3 shows the respective Kaplan-Meyer plots for few genes that showed significant differences in survival between patients belonging to peaks 1 and 2. Supplementary Figure 4 shows the Kaplan-Meyer plot for all 96 genes associated with prognosis.

**Figure 3.**
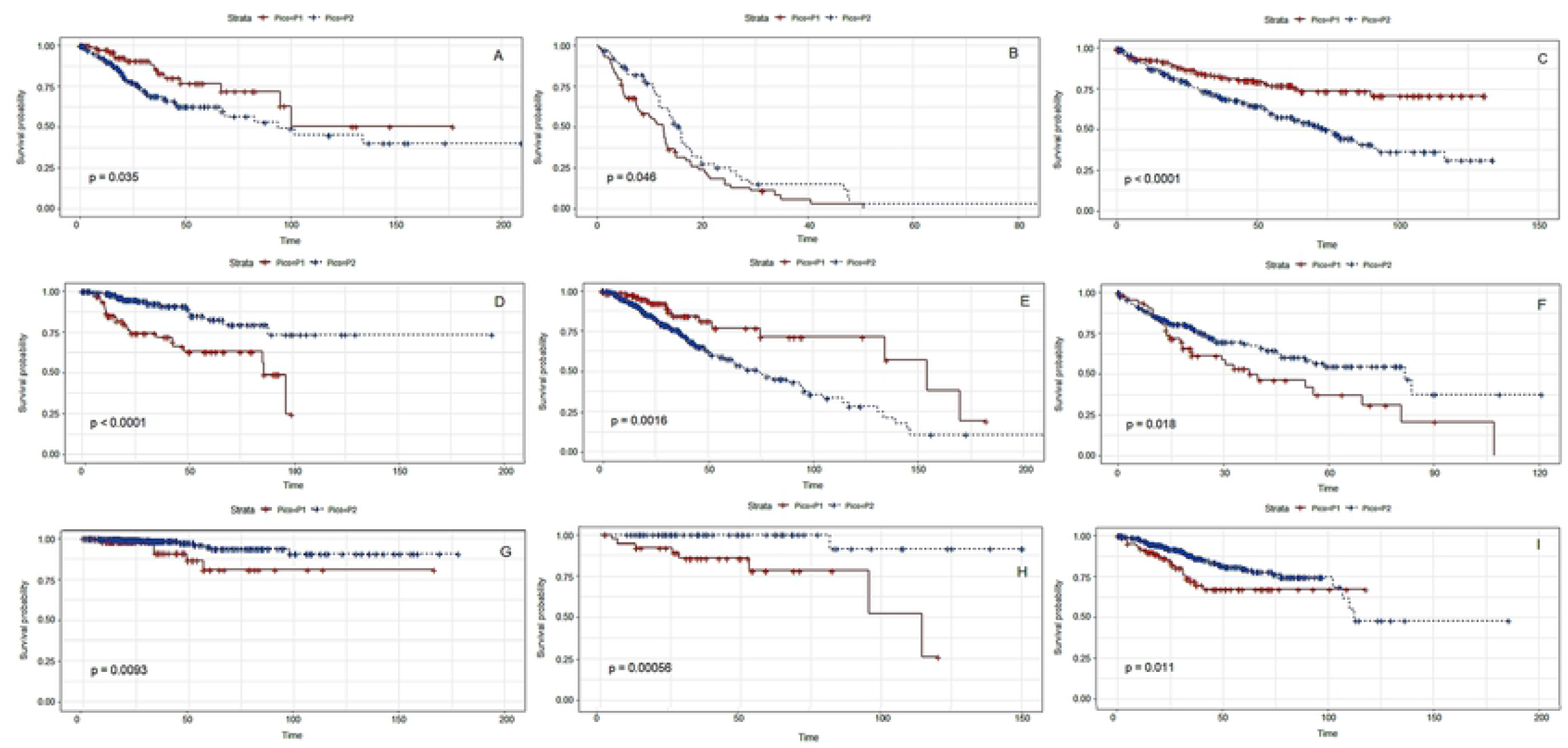
Kaplan-Meier plots of representative genes for each tumor type (one gene per tumor, arbitrarily selected). P1 and P2 correspond to the two modes of the bimodal distribution. A) Gene ZNF304 for CESC; B) Gene ZBTB45 for GBM; C) Gene XRRA1 for KIRC; D) Gene DYNC2LI1 for KIRP; E) Gene CDC42 for LGG; F) Gene KDM5D for LIHC; G) Gene ANXA1 for THCA; H) Gene LAIR1 for THYM and I) Gene HNF1B for UCEC.

## Discussion

A new genome-wide method is presented to identify genes with bimodal patterns of expression using GMM analysis [14] for the stratification of samples. GMM has been previously used in the analysis of gene expression data [23–25] but to our knowledge this is the first application of such a method for the identification of genes with bimodal expression patterns.

The applicability of the method is shown by using gene expression and clinical data for 25 tumor types available from TCGA. We identified 554 genes unique with bimodal gene expression (Table 1). Forty-six of them were identified as bimodal in more than one tumor type. Several of them have been reported previously as having a bimodal expression pattern. One of them, ERAP2, had been found by [4] to have bimodal gene expression in human skeletal muscle. The same report from [4] found that GSTM1 has a bimodal expression pattern in muscle tissue. Other genes include RPS27, found by [11] to have bimodal expression in several tumor types, and USP9AY, found to show bimodality in endometrium [26]. Interestingly, among the 46 genes with bimodality in more than one tumor type, 12 are mapped to the Y chromosome (p<10^-5^), an unexpected observation due to the low gene density in this chromosome. As reviewed by [27], some genes on the Y chromosome have dosage-sensitive functions, which might be related to a bimodal expression pattern. This remains to be further explored.

Ninety-six, out of 554 genes with bimodal gene expression in all tumor types analyzed here, were identified as having differential prognosis when patients belonging to the two different modes were compared. Expression of several genes identified by us are known predictors of clinical outcome in different tumor types including ANXA1 [28], FOXJ3 [29, 30] and CDC25 [31, 32], among many others. However, the great majority of these reports only associate expression with prognosis. On the other hand, we associate the bimodal expression pattern with prognosis. To our knowledge, only [11] have associated the bimodal expression pattern of RPS27 with clinical outcome in several tumor types, a gene also observed in our data. This makes our analysis the first one, to our knowledge, to explore the association between the modes of gene expression distribution with prognosis in a genome-wide context.

Several issues should be considered in the interpretation of our results. For example, cellular heterogeneity within samples in a given cohort is a factor that can generate genes with bimodal expression. In our case, this is minimized by the fact that TCGA samples are selected for high tumor cell content but this issue should be critically considered when more heterogeneous cohorts are analyzed. Furthermore, clinical and/or biological features should be considered when interpreting data from our method. For example, in cancer studies one should be careful with cohort heterogeneity regarding staging and progression, among many other clinical features.

We envisage that our method will be a useful tool for the genome-wide identification of genes with bimodal pattern of expression. The computational pipeline to execute the method is freely available at https://github.com/LabBiosystemUFRN/Bimodality_Genes.

---

**Supp. Figure 1.**
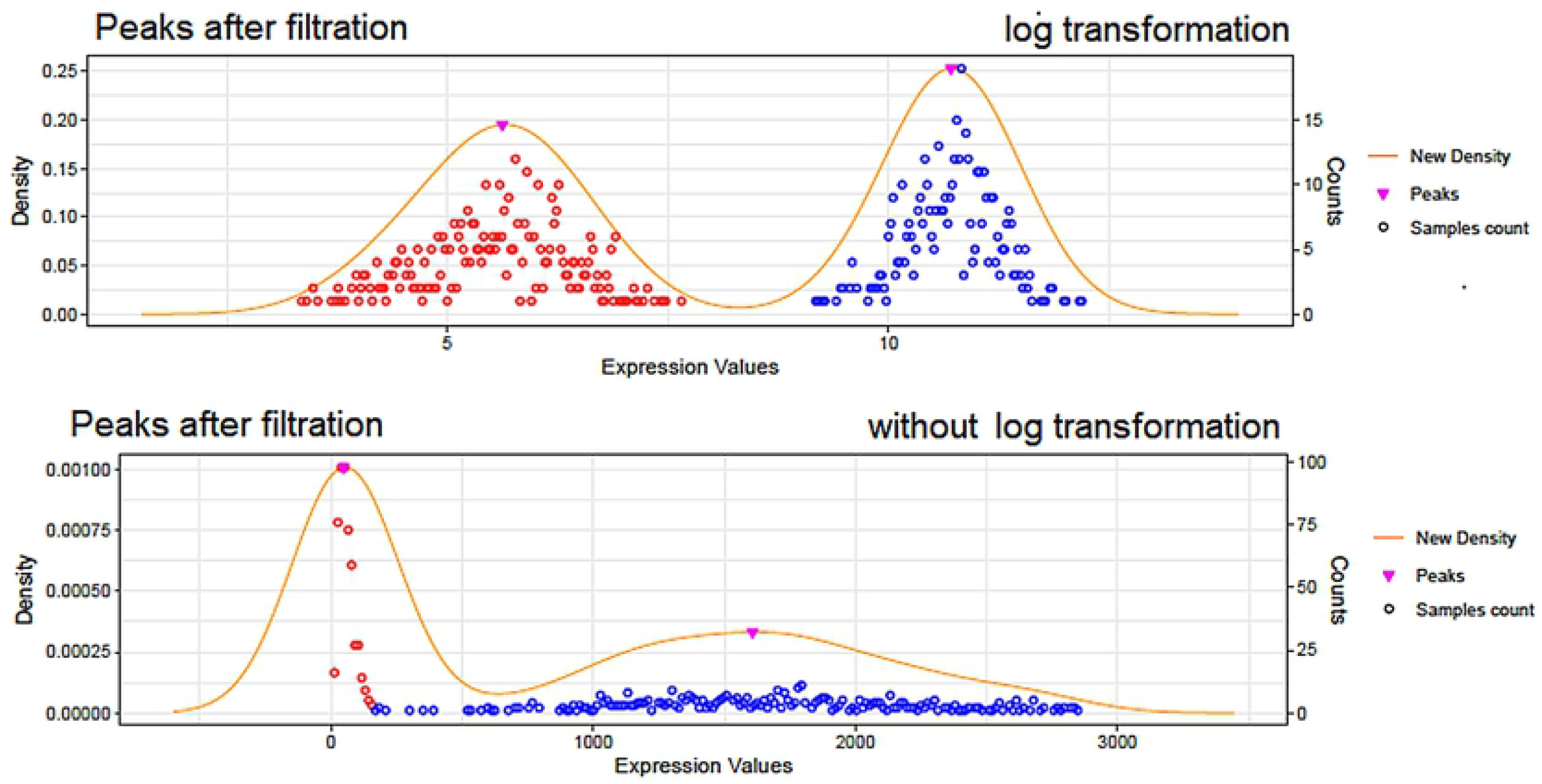

**Supp. Figure 2.**
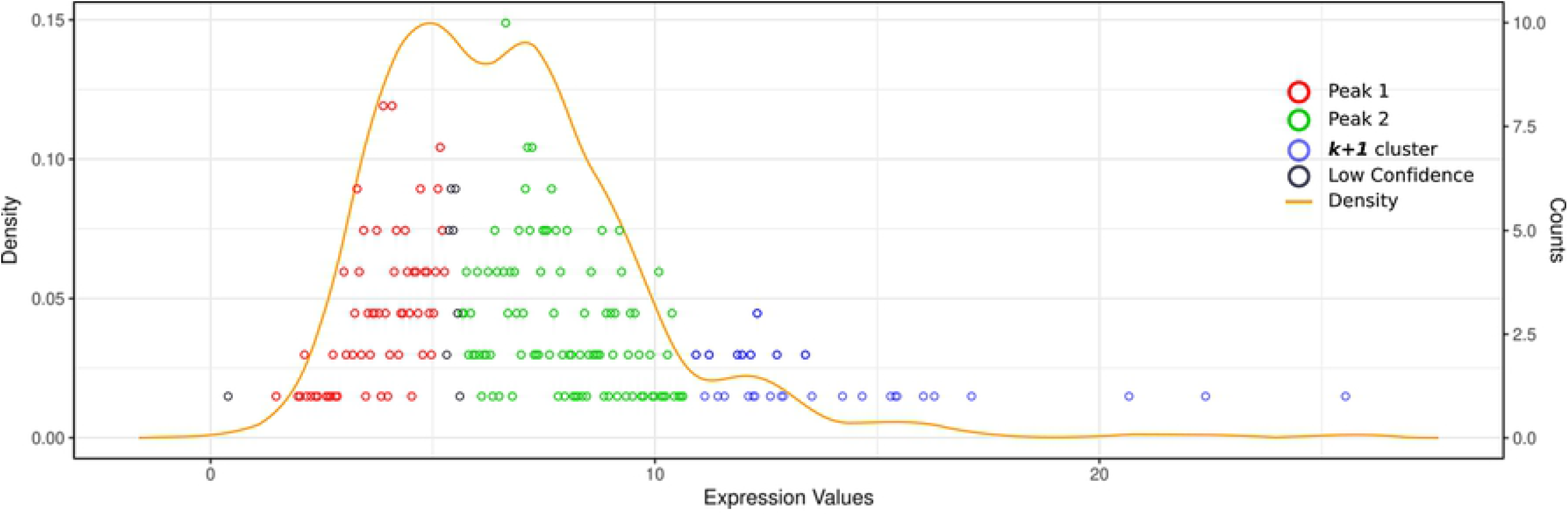

**Supp. Figure 3.**
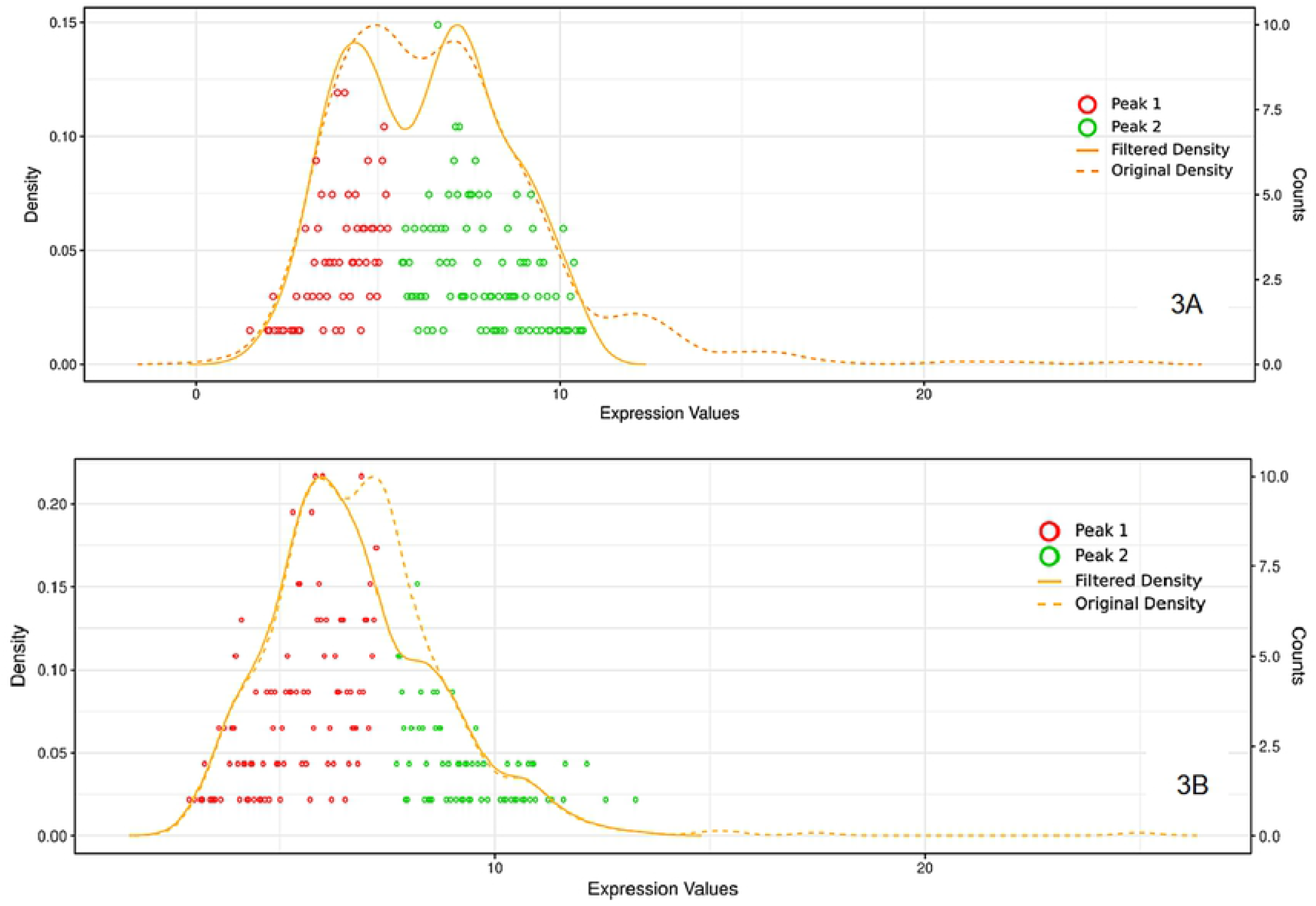

**Supp. Figure 4.**
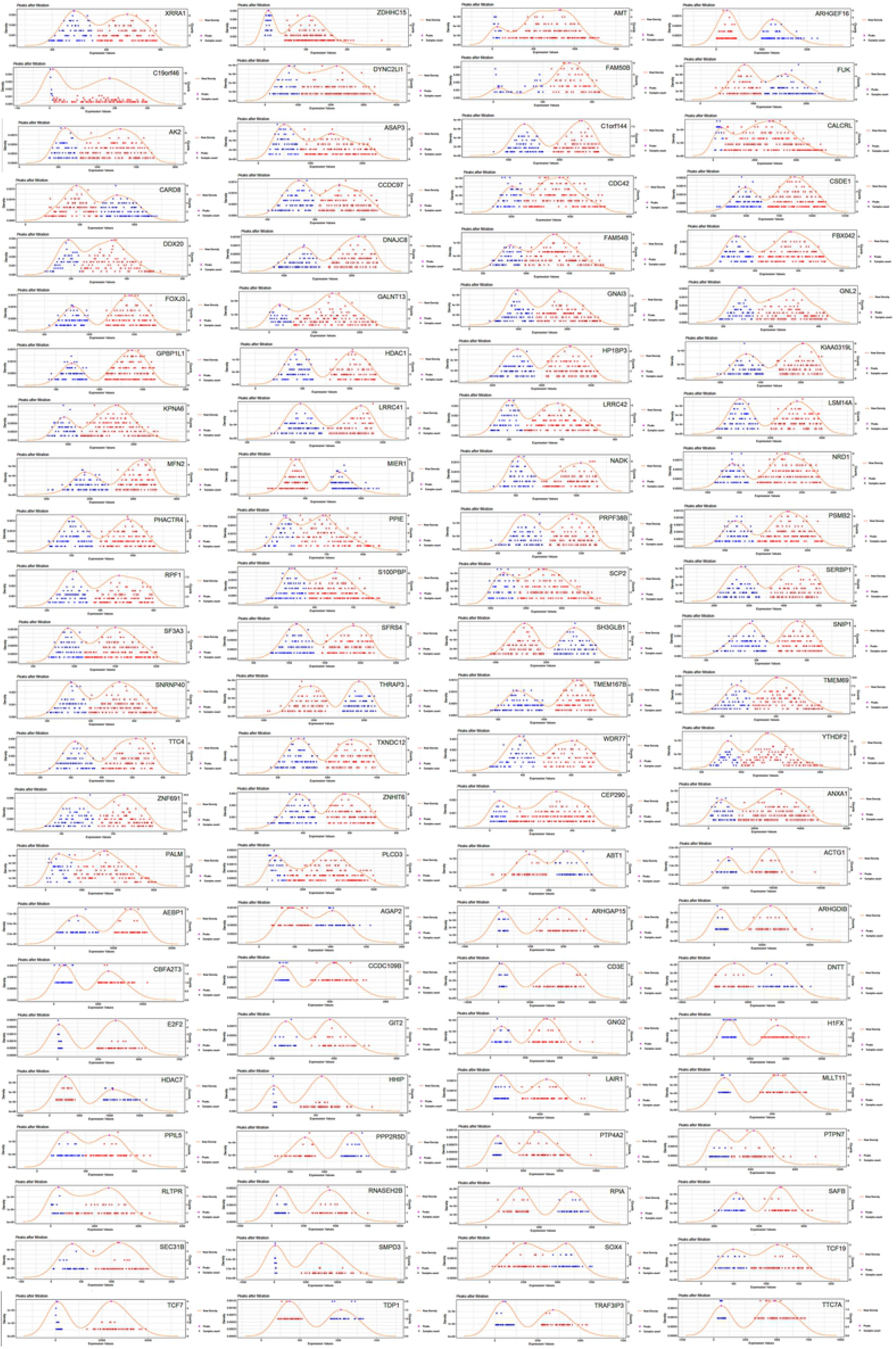

**Supp. Figure 5.**
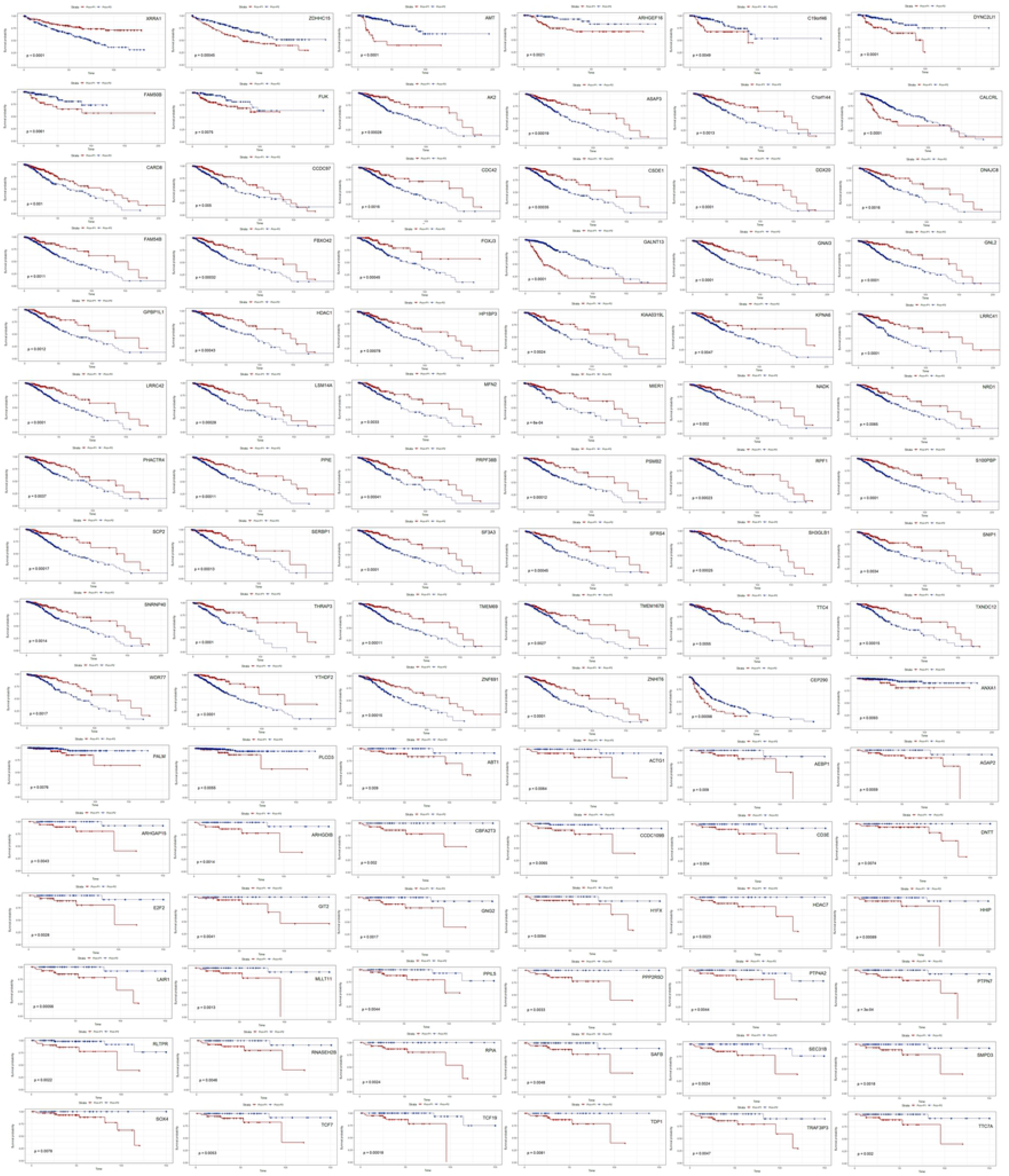

